# MondoA Drives Malignancy in cALL through Enhanced Adaptation to Metabolic Stress

**DOI:** 10.1101/2020.11.01.363986

**Authors:** Alexandra Sipol, Erik Hameister, Busheng Xue, Andreas Petry, Carolin Prexler, Maxim Barenboim, Rebeca Alba Rubio, Michaela C. Baldauf, Davide G. Franchina, Juliane Schmaeh, Guenther Richter, Gunnar Cario, Thomas G.P. Grünewald, Agnes Görlach, Dirk Brenner, Jürgen Ruland, Poul H. Sorensen, Stefan E.G. Burdach

**Author notes:** Corresponding author: Stefan Burdach, MD, PhD, Children’s Cancer Research Center and Department of Pediatrics, Technische Universität München (TUM), D-81664 München fon or -2261, mailto. Authors contributed equally (Alexandra Sipol, Erik Hameister). Shared senior authorship (Poul H. Sorensen, Stefan E.G. Burdach). MLL Münchner Leukämielabor GmbH Max-Lebsche-Platz 31, 81377 München, mailto.

## Abstract

Cancer cells are in most instances characterized by rapid proliferation and uncontrolled cell division. Hence, they must adapt to proliferation-induced metabolic stress through intrinsic or acquired anti-metabolic stress responses to maintain homeostasis and survival. One mechanism to achieve this is to reprogram gene expression in a metabolism-dependent manner. MondoA (also known as MLXIP), a member of the MYC interactome, has been described as an example of such a metabolic sensor. However, the role of MondoA in malignancy is not fully understood and the underlying mechanism in metabolic responses remains elusive. By assessing patient data sets we found that MondoA overexpression is associated with a worse survival in pediatric common acute lymphoblastic leukemia (cALL). Using CRISPR/Cas9 and RNA interference approaches, we observed that MondoA depletion reduces transformational capacity of cALL cells *in vitro* and dramatically inhibits malignant potential in an *in vivo* mouse model. Interestingly, reduced expression of MondoA in patient data sets correlated with enrichment in metabolic pathways. The loss of MondoA correlated with increased tricarboxylic acid (TCA) cycle activity. Mechanistically, MondoA senses metabolic stress in cALL cells by restricting oxidative phosphorylation through reduced PDH activity. Glutamine starvation conditions greatly enhance this effect and highlight the inability to mitigate metabolic stress upon loss of MondoA in cALL. Our findings give a novel insight into the function of MondoA in pediatric cALL and support the notion that MondoA inhibition in this entity offers a therapeutic opportunity and should be further explored.

**Key Points:** MondoA maintains aggressiveness and leukemic burden in common ALL, modulating metabolic stress response.

## Introduction

Acute lymphoblastic leukemia (ALL) is the most frequent childhood malignancy, with common B-precursor ALL (cALL) being its most common subgroup. In spite of major advances in treating ALL, it remains a leading cause of cancer death in children, and relapse is associated with a particular poor outcome^1^. Additionally, current treatment regimens result in considerable long-term toxicity^2,3^. Even patients that have been treated successfully face a reduced life expectancy with a 13% cumulative incidence of death from any cause including leukemia recurrence, subsequent second cancers, or a risk of dying from cardiac-related events^4^. This clearly highlights the need for a better understanding of this malignant entity in initial treatment as well as reducing toxicity.

Almost a century ago, Otto Warburg observed that tumor cells generate lactate through glycolysis even in the presences of oxygen^5^. It is thought that intermediaries of the glycolytic pathway are used for *de novo* synthesis of biomolecules such as fatty acids, nucleic acids and amino acids^6^. Furthermore, tumor cells must adapt to proliferation-induced metabolic stress, and require anti-metabolic stress responses to maintain homeostasis and survival^7^. Cells reprogram their gene expression in a metabolism-dependent manner to support their proliferation. One such mechanism is driven by energy sensing molecules which can drive cellular energy homeostasis in response to starvation conditions – AMP-activated protein kinase (AMPK) and hypoxia-inducible factor 1-alpha (HIF1α) are well known examples^8,9^. Another protein that has been proposed to influence energy homeostasis by sensing intracellular energy states is MAX (Myc-Associated factor X)-Like protein X (MLX) Interacting Protein (MLXIP), also known as MondoA^10-14^.

MondoA acts as a pro-glycolytic basic helix-loop-helix leucine zipper transcription factor and is part of the highly complex MYC interactome, which comprises multiple transcription factors (MYC/MAX/MAD/MLX/MondoA). The key players of this network, MYC, MAD and MondoA, differentially mediate proliferation, differentiation or metabolism by competing for heterodimerization with limited interactors, resulting in varying signaling outcomes. Hence, the MYC interactome can regulate both proliferation as well as the necessary adaptation to proliferative and metabolic stress^15,16^. For example, in glucose dependent tumor entities, such as triple negative breast cancer or malignant melanoma, MYC and MondoA behave antagonistically^17,18^. In contrast, in a glutamine dependent malignancy, neuroblastoma, these two proteins cooperate to confer malignancy^16^. We also previously showed that MondoA is required for the proliferation and stemness features of cALL^19^. MondoA is known to be a potent negative regulator of glucose uptake and is highly sensitive to increasing intracellular glucose-6-phosphate (G6P) levels allowing for adaptive transcriptional responses to changes in extracellular glucose concentrations^15,16^. Here we report that MondoA is overexpressed in cALL and correlates with relapse and decreased survival. We show that depletion of MondoA results in loss of proliferative capacity and features of malignancy. Interestingly, the loss of MondoA leads to increased oxidative phosphorylation. Mechanistically, MondoA mediates malignancy by facilitating adaptation to metabolic stress by restricting TCA cycle activity. MondoA reduces PDH activity in glutamine scarce environments through increased PDK expression. We therefore hypothesize that MondoA mediates malignancy in cALL by functioning as a metabolic stress sensor in pediatric cALL. This potentially warrants the inhibition of MondoA as a novel therapeutic target in patients with cALL.

## Materials and Methods

### Cell lines

Human cALL cell lines 697, REH and Nalm6 were obtained from the German Collection of Microorganisms and Cell Cultures (DSMZ). Cell lines were checked routinely for purity (surface antigens, HLA-phenotype and STR authentication via DSMZ online STR analysis tool) and Mycoplasma contamination.

### Primary patient-derived material

Primary human samples were obtained with ethics approval of the Institutional Review Board (approval ID: 2562/09). Pre-therapy cALL samples were obtained from bone marrow (n=61) or peripheral blood (n=11) of children treated according to the ALL-BFM study of the Society for Pediatric Oncology and Hematology (GPOH). All patients provided written informed consent. Mononucleated cells (MNC) from bone marrow or peripheral blood were processed by gradient centrifugation (Biocoll separation solution, Biochrom) and cryopreserved.

### Generation of MondoA knockdown clones

For generation of an inducible MondoA knock down (MKD), 697, REH, Nalm6 cALL cells were infected with lentivirus (MOI: 1:10) containing a pTRIPZ vector with either an shRNA against MondoA (clone V2THS_96171 sequence 5’-TGAAACATGCCCAGCAGGG-3’, clone V2THS_96169 sequence 5’-TCCCTGAGCAGTTGTTCTG-3’; Dharmacon) or respective non-targeting control shRNA. Single-cell cloned 697, REH, and Nalm6 infectants were selected in 0.5 µg/ml Puromycin (Invitrogen). MKD efficacy upon doxycycline-treatment (1.0 µg/ml) was confirmed by real-time PCR and western blot (WB). For lentivirus production 5.5 × 10^6^ HEK293T cells were seeded one day before transfection and cultured in DMEM (Invitrogen) supplemented with 10% FBS, 100 units/ml penicillin and 100 µg/ml streptomycin (both Invitrogen). Supernatant was harvested 48h after transfection and lentiviral particles were isolated by filtration.

### Generation of MondoA knock out clones

We designed our sgRNA (Methabion) to target the Cas9 nuclease to exon 10 of MondoA to ensure that the encoded mutant proteins were dysfunctional and that all known splice variants would be affected. The design was performed using the Brunello Database that identifies optimal target sequences with a minimum of potential off-target sites^20^. Oligonucleotides were cloned into lentiviral vector LentiGuide-Puro (Addgene plasmid #52963). Lentiviral constructs LentiCas9-Blast (Addgene Plasmid #52962) and LentiGuide-Puro plasmid (Addgene plasmid #52963) with ligated selected sgRNA were transfected into HEK293T. Viral supernatants were isolated 48h after transfection and filtered. Nalm6 cell line was transduced with viral supernatant. Blasticidin-resistant cells were subsequently transduced with viral supernatant containing LentiGuide-Puro plasmid (Addgene plasmid #52963) with ligated selected sgRNA. Stable infectants were selected in 2 µg/mL puromycin (pSIREN-RetroQ). Limiting dilutions were performed to isolate single cells in 96-well plates. Mutants and control clones were identified by WB. MondoA protein levels were measured using an anti-MondoA rabbit monoclonal antibody (Proteintech; 13614-1-AP) diluted to 1:500 in 5% skim milk (Sigma-Aldrich) in Tris buffered saline solution containing Tween 20 (Sigma-Aldrich).

### Generation of MondoA overexpressing clones

Viral supernatant with Lenti ORF particles with MondoA (Myc-DDK tagged) construct for overexpression was purchased from (Origen CAT#: RC222157L3V). Cas9-transduced Nalm6 cells were transduced. Stable infectants were isolated after selection in 2 µg/mL Puromycin.

### Microarray analysis

RNA was amplified and labeled using Affymetrix GeneChip Whole Transcript Sense Target Labeling Kit. cRNA was hybridized to Affymetrix Human Gene 1.0 ST arrays and analyzed by Affymetrix software expression console, version 1.1. For the data analysis, robust multichip average (RMA) normalization was performed, background correlation, quantile normalization, and median polish summary method. Microarray data were deposited at the gene expression omnibus (GSE76277). Analysis was performed with signal intensities that were log2-transformed for equal representation of over- and under-expressed genes. Gene-set enrichment analysis (GSEA) and set-to-set pathway analyses of leading-edge genes were conducted with the GSEA tool (http://www.broad.mit.edu/gsea) in default parameters^21-23^. Comparison of independent published microarray studies was performed as previously described^24^. Briefly, microarray data, which were all generated on Affymetrix HG-U133Plus2.0 chips, were downloaded from the Gene Expression Omnibus (GEO) or the EBI and normalized simultaneously with RMA and brainarray CDF files (v19, ENTREZG) yielding one optimized probe-set per gene. Accession codes are given in supplemental data^25^.

### Metabolism Assay

OCR was analyzed on a XF96 Extracellular Flux Analyzer (Seahorse Bioscience). Cells were plated in nonbuffered RPMI 1640 media with 10 mM glucose. Measure^-^ ments were obtained under basal conditions and after the addition of 1 mM oligomycin, 1mM FCCP and 1mM Rotenone / Antimycin A.

### Measurement of intracellular ROS

Cells were exposed to 30 min incubation with 5 mM dichlorofluorescein diacetate (DCF-DA, Sigma). Cells were analyzed by flow cytometry.

### Mice and *in vivo* experiments

Rag2^-/-^*γc* ^-/-^ mice on a BALB/c background were obtained from the Central Institute for Experimental Animals (Kawasaki) and maintained under pathogen-free conditions and approval of the local authorities. Experiments were performed in (6-16)-week-old female mice, n=5 for each experimental group. For *in vivo* tumor growth 5 × 10^6^ leukemic cells in 200 µl PBS were injected into the tail veins of mice, which received doxycycline containing food (Altromin). Five weeks after intravenous injection, mice were euthanized and leukemic burden was monitored by analysis of human CD10 expression in blood, spleen and bone marrow. Liver and spleen of treated mice were formalin-fixed for histology. In experiment with MKO Nalm6 cell lines, 8 mice were injected with 5 × 10^6^ MKO Nalm6 cells and 9 mice with Cas9 CTRL Nalm6 cells. These mice received standard no doxycycline containing food.

## Results

### MondoA is overexpressed in primary cALL and correlates with higher relapse and decreased survival in cALL subsets

Previously we reported that MondoA is overexpressed in cALL in comparison to 36 normal adult and fetal tissues^19^. To further investigate the gene expression pattern of MondoA in different human cancers, we compared microarray datasets from publicly available data sources (Figure 1A)^25^. We analyzed 1042 independent cancer samples, including 533 pediatric tumors, consisting of a wide array of tumor entities including 33 samples of cALL. MondoA expression was observed to be 2-fold higher (median expression intensity (MEI) = 395.7, SD = 280.6) relative to all other pediatric tumors in the data set (MEI = 148.5, SD = 42, p<0.001) (Figure 1A). Looking more closely at hematopoietic neoplasms in the same data set, MondoA was also most highly expressed (MEI = 395.7, SD = 280.6) in pediatric cALL compared to any other hematopoietic malignancy (MEI average = 152.42, SD = 32.48, p<0.001) (Figure 1B). Another publicly available data set using RNA-Seq analysis (UCSC Xena) identified pediatric cALL as having the highest MondoA expression (Median normalized count (MNC) = 15.36, SD = 1.06) compared to all other pediatric tumors (MNC = 12.35, SD = 0.56, p<0.001) (Figure 1C). We therefore validated the MondoA expression in pediatric cALL in our own patient population using an independent method. From eleven patients with cALL, we derived peripheral blood mononuclear cells (PBMC) and performed quantitative RT-PCR (qRT-PCR). Three blood samples from healthy donors were used as controls. cALL bearing patients had a significantly increased expression level of MondoA compared to the healthy controls (p = 0.0499) (Figure 1D). We therefore asked if prognostic information could be drawn from an altered MondoA expression. To address this question, we analyzed MondoA mRNA expression in 61 bone marrow samples from the BFM (Berlin, Frankfurt, Münster) study group patients with primary cALL. All 61 samples were negative for other known prognostically relevant molecular markers, including TEL/AML1, BCR/ABL, and MLL/AF4. Samples were either grouped into a very high-risk group (very HR) (n = 31), defined as minimal residual disease (MRD) ≥10^−3^ after induction treatment, or a non-high-risk group, defined by prednisone good response, remission at day 33, MRD measurable at <10^−3^ on treatment day 33 and at week 12 (n = 30). MondoA expression was 1.63-fold higher in the very HR patient set (p = 0.0294) (Figure 1E). Analyzing publicly available survival data (UCSC Xena) of pediatric patients with BCR-ABL negative ALL demonstrated also that high MondoA expression in primary bone marrow samples corresponded with a significantly decreased survival (p = 0.03), cohort of patients with minimal residual disease on day +29 of treatment protocol (n=67, median age 14 y, TARGET) (Figure 1F). Initially, we conclude that MondoA is greater expressed in patients with pediatric cALL compared to other pediatric malignancies, and that MondoA overexpression is associated with a worse patient outcome in cALL.

**Figure 1.**
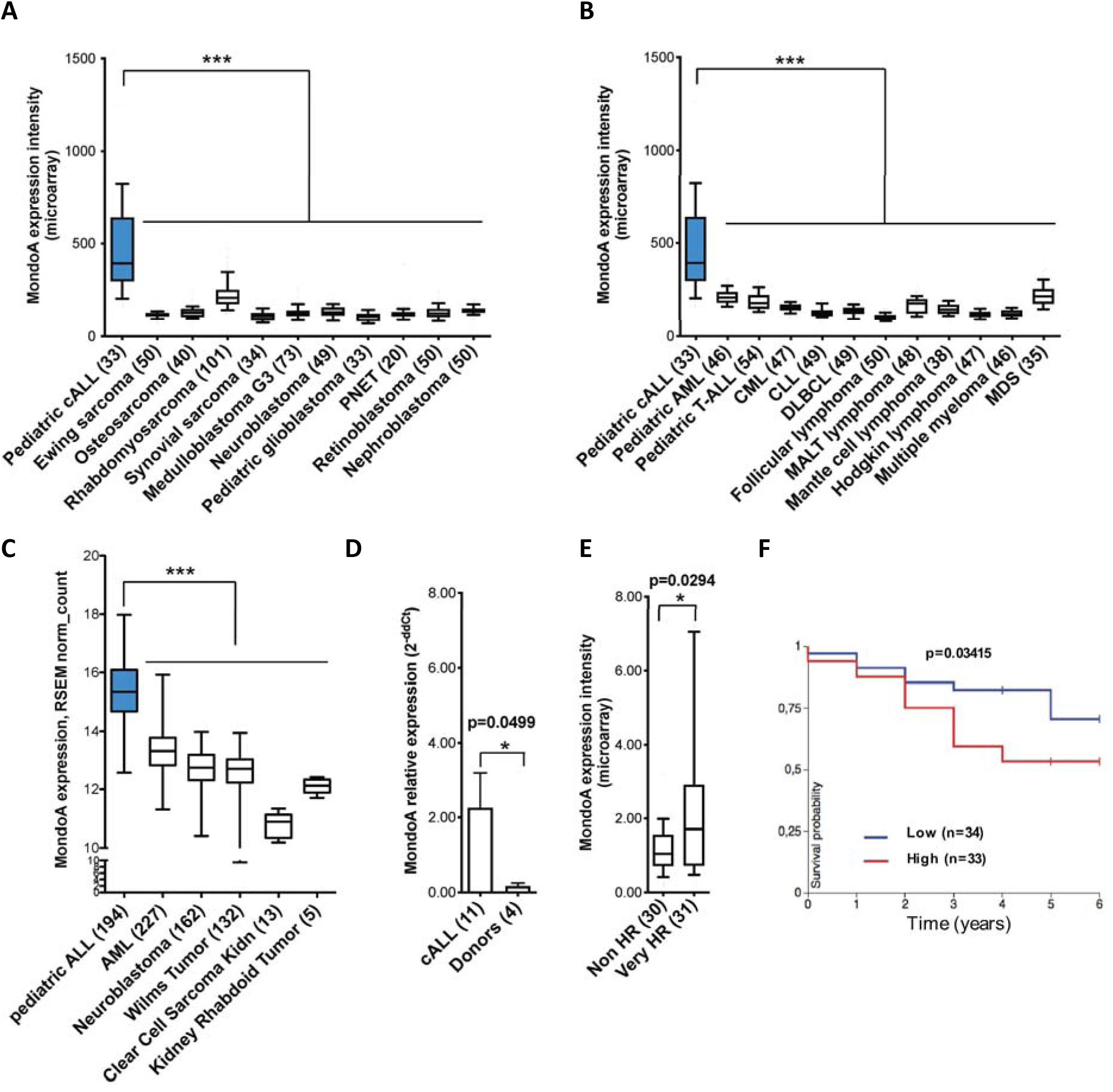
MondoA is highly expressed in primary cALL and correlates with relapse risk. **(A)** MondoA (ID 22877_at) expression in pediatric cALL relative to pediatric solid tumors (n=533, p<0.001, one-way ANOVA with Bonferroni’s Multiple Comparison Test). cALL, pediatric common B-precursor acute lymphoblastic leukemia; PNET, primitive neuroectodermal tumor. Numbers of analyzed samples shown in brackets. All datasets were normalized simultaneously using RMA and custom microarray (v15 ENTREZG) CDF files. Data depicted as box-plots. Whiskers indicate the 10^th^ and 90^th^ percentile. Data presented in linear scale. MondoA Microarray ENTREZG probeset ID is 22877_at. **(B)** MondoA (ID 22877_at) expression in lymphoid and myeloid neoplasms (n=542, p<0.001, one-way ANOVA with Bonferroni’s Multiple Comparison Test). AML, acute myeloid leukemia; T-ALL, T-cell acute lymphoblastic leukemia; CML, chronic myeloid Leukemia; CLL, chronic lymphocytic leukemia; DL-BCL, diffuse large B-cell lymphoma; MALT lymphoma, mucosa associated lymphoid tissue lymphoma; MDS, myelodysplastic syndrome. **(C)** MondoA expression in pediatric cALL relative to AML and other paediatric tumors (RNAseq data, UCSC Xena, n=733, p<0.001, one-way ANOVA with Bonferroni’s Multiple Comparison Test). **(D)** MondoA relative expression by qRT-PCR in PBMC samples from patients (n=11) with primary pediatric cALL compared to three PBMCs from healthy donors (n=4), normalized to donor PBMC sample. Results of two independent experiments in duplicate are presented as means ± SEM (p=0.0499, Welch’s t test). **(E)** MondoA overexpression correlates with relapse risk in cALL. MondoA expression 63% higher (p=0.0294, Welch’s t test) in very high-risk group (very HR) (n=31) as compared to non-high-risk group (Non HR) (n=30), defined by prednisone good response (PGR), remission on day 33 of induction therapy, MRD-MR or MRD-SR. All cases are negative for prognostically relevant molecular markers TEL/AML1, BCR/ABL, MLL/AF4 (data from the BFM study group). Bars indicate median, boxes represent middle 50% of data. Whiskers indicate 10^th^ and 90^th^ percentile. **(F)** Kaplan-Meier curve indicating significant difference in survival of cALL patients with positive MRD on day +28 depending on MondoA expression. All cases are negative for BCR/ABL (n=67, p=0.03415).

### MondoA silencing reduces proliferation and clonogenicity *in vitro*, and aggressiveness *in vivo*

To investigate the potential functional role of MondoA in cALL, we employed RNA interference and CRISPR/Cas9 mediated gene editing (Cas9) to test the *in vitro* effects of MondoA-suppression on cALL proliferation and clonogenicity. To this end, we generated a doxycycline-inducible short hairpin RNA (shRNA) expression system in three independent cALL cell lines (Nalm6, 697 and REH). MondoA knockdown (MKD) efficiency of shRNAs was confirmed by PCR qRT-PCR and Western blot (WB) (Figures 2A, B). In a colony formation assay we seeded MondoA silenced and control cells at low density into methylcellulose containing doxycycline. We observed a significant two-fold decrease of colony numbers after 14 d in the experimental conditions used in Naml6 and 697 cell lines (p<0.01) (Figure 2C, Suppl. figure S1A, B). We observed a significant reduction in proliferation in Nalm6, REH and 697 cell lines upon MKD in a BrdU proliferation assay (p<0.001 at 48 hour and 72 hour) (Figure 2 D, E, F).

**Figure 2.**
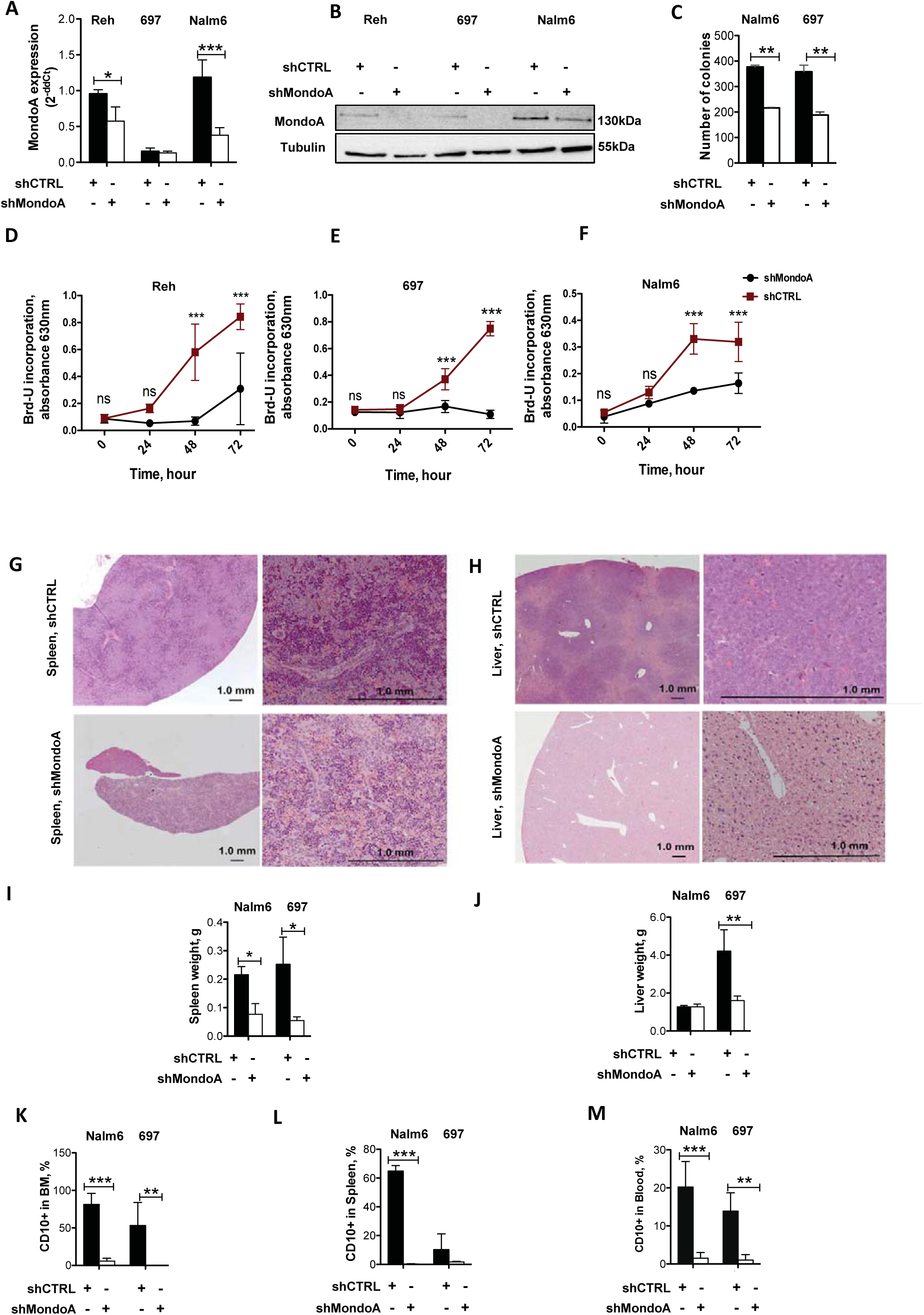
Knockdown of MondoA (MKD) reduces proliferation and clonogenicity of cALL cells *in vitro* and tumorigenicity *in vivo*. **(A)** Knockdown efficiency of MondoA by doxycycline inducible shRNA against MondoA compared to control shRNA (shCTRL, Tet-On) by qRT-PCR after treatment with doxycycline for 48h. Data are mean and SEM of n=6 experiments, p<0.01 (Reh) and p<0.001(Nalm6). **(B)** Western blot of knockdown efficiency after 48h of doxycy^-^ cline. **(C)** Contact-independent growth in Nalm6 and 697 cells stably transfected with doxycycline inducible shRNA either against MondoA or control. Data are mean of n=2 experiments, p<0.01, Mann Whitney test. **(D, E, F)** Long-term proliferation measured by Brd-U assay. Control and MKD cells pretreated with doxycycline for 48h (n=6 for each time point, p<0.001, 2way ANOVA). **(G, H)** Xenograft transplantation of Nalm6 and 697 cells stably transfected with doxycycline inducible shRNA infectants in Rag2^−/−^yc^−/−^ mice. H&E staining of formalin-fixed paraffin-embedded liver and spleen (scale bar = 1.0 mm). Images captured from hematoxylin-and-eosin staining of the 4µm sections of paraffin blocks by Axioplan2 microscopy (Carl Zeiss, 1997) and AxioVs40 V.4.8.2.0 software. **(I)** Weight of spleen in g (grams). Cell lines either expressing shRNA against MondoA or control, as indicated (n=5 in each group, p<0.05). **(J)** Weight of liver in g. Cell lines either expressing shRNA against MondoA or control, as indicated (n=5 in each group, p<0.01, results are presented as mean ± SEM; Mann Whitney test). CD10^+^ leukemic blasts in bone marrow **(K)**, spleens **(L)**, and blood **(M)** in cell lines either expressing shRNA against MondoA or control, as indicated (p<0.001 (Nalm6) and p<0.01 (697), results are presented as mean ± SEM; Mann Whitney test).

To extend these findings to an *in vivo* setting, MKD clones generated initially for *in vitro* experiments and control Nalm6 and 697 cells were injected into the tail veins of immune compromised mice. Serial sectioning of liver and spleen tissue, including measurements of liver and spleen weights as well as the relative amount of human CD10^+^ leukemic blasts in blood, bone marrow and spleen were then assessed. The reduced number of leukemic cells was also confirmed by hematoxylin and eosin staining of formalin-fixed paraffin-embedded spleen and liver tissue (Figure 2G, H). Knockdown of MondoA was associated with a significant reduction in average size and weights of spleen and liver (p<0.05 and p<0.01) (Figure 2 I, J, S1C). A significant decrease of CD10^+^ leukemic blasts in bone marrow, spleen and blood for Nalm6 cells as well as in bone marrow and blood for 697 cells was also observed (p<0.001 (Nalm6) and p<0.01 (697)) (Figure 2K, L, M).

Since shRNA mediated knockdown did not fully suppress MondoA expression as evidenced by qRT-PCR (Figure 2A), validation through a CRISPR/Cas9 approach to generate an isogeneic Nalm6 cALL cell lines with a complete knockout of MondoA were generated. Following stable Cas9 MondoA knockout (MKO), MondoA was undetectable at the protein level (Figure 3A). In line with the MKD experiments, MKO cells showed reduced proliferation and formed fewer colonies in comparison to control cells (Figure 3B, C, D, S1D). Consistently, the tumorigenic potential of xenotransplanted MondoA deficient Nalm6 cells was significantly decreased *in vivo*. The survival of mice was monitored for 56 days at which time all 9 mice injected with control Nalm6 cells had to be sacrificed (Figure 3E, p<0.001). In sharp contrast, mice injected with MondoA deficient Nalm6 cells showed no sign of disease. Mice in the control group showed significant spleen enlargement and hepatomegaly in post-mortem analysis compared to the experimental group (p<0.001 and p<0.01) (Figure 3F, G, S1E). Furthermore, detectable CD10^+^ positive blasts by flow cytometry was below 1 % in bone marrow, spleen and liver of mice initially injected with MKO Nalm6 cells (data not shown). To summarize, our data suggests that stable MondoA knockdown or knockout decreases proliferation, clonogenicity and aggressiveness of cALL cells *in vitro* and *in vivo*.

**Figure 3.**
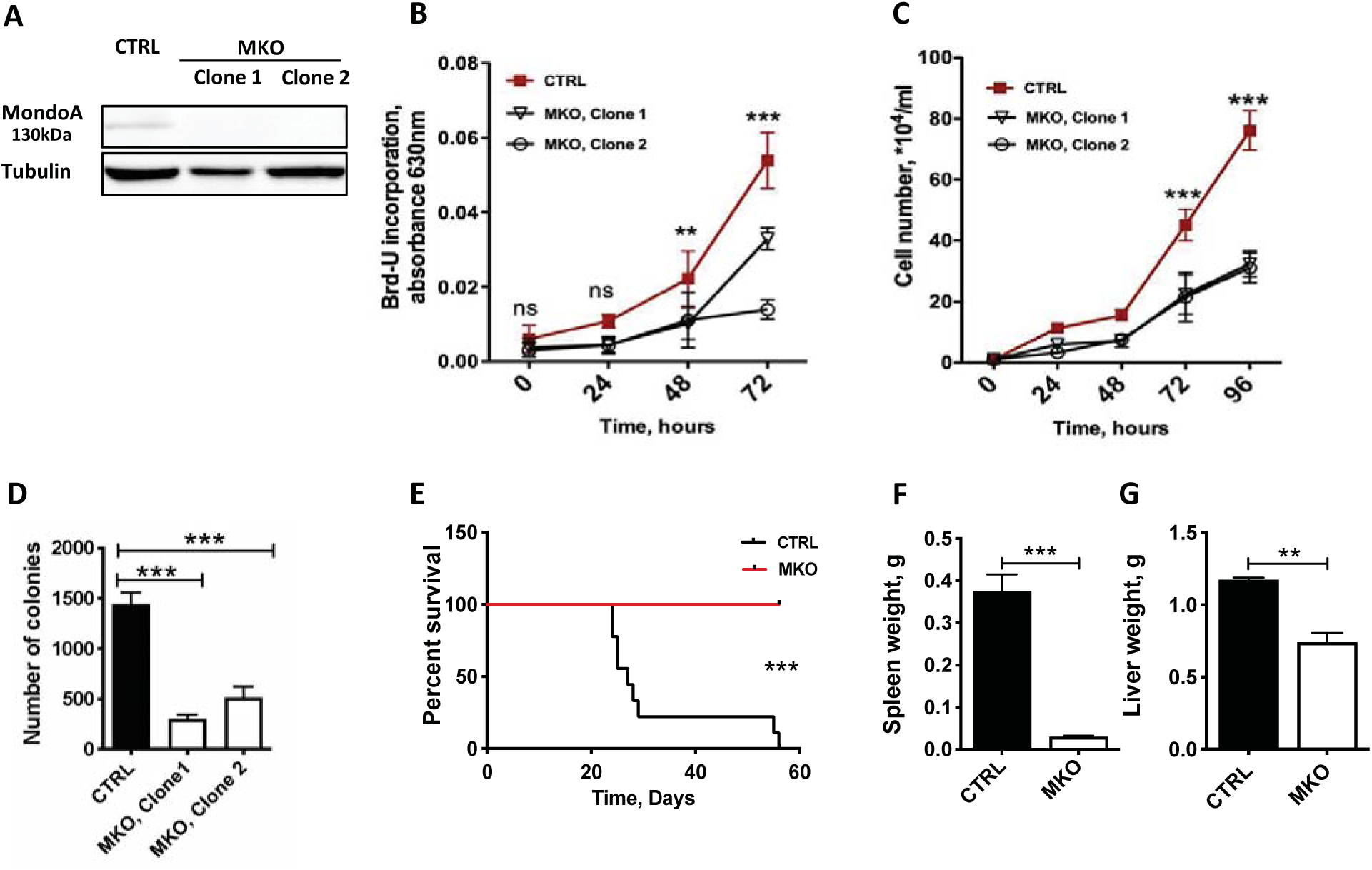
CRISPR/Cas9 mediated MondoA Knockout (MKO) reduces proliferation and clonogenicity of cALL cells *in vitro* and tumorigenicity *in vivo*. **(A)** Representative Western blot with anti-MondoA antibody of Nalm6 MKO clone 1 and clone 2 lysates; Cas9-only transfected Nalm6 cALL cell line as control; alpha-tubulin staining as loading control. **(B)** Long-term proliferation measured with Brd-U assay (n=6 for each time point, p<0.01 at 48h and p<0.001 at 72h, 2way ANOVA). **(C)** Long-term proliferation measured by direct cell counting. Cas9-only transfected Nalm6 cALL cell line and with sgRNA against MondoA, were grown under normoxia (20% O_2_) (n=6 for each time point, p<0.001 at 72h and 96h, 2way ANOVA). **(D)** Contact independent growth of Nalm6 cells with CRISPR/Cas9 mediated MKO. Quantitative evaluation in methylcellulose-based colony formation assay (n=4, p<0.001, results are presented as mean ±SD; one-way ANOVA). **(G)** Survival curves of mice transplanted with Nalm6 knockout for MondoA and Cas9 only control cells (n=8 for MKO group, n=9 for CTRL, p<0.001). **(F, G)** Xenograft transplantation of Nalm6 cells with either MKO (n=8) or control (n=9) in Rag2^−/−^yc^−/−^ mice. Weight of liver and spleens in g (grams) (p=0,0026 (liver) and p=0.0008 (spleen), results are presented as mean ± SD; Unpaired t test).

### MondoA suppresses MYC-targets, fatty acid metabolism, oxidative phosphorylation and ROS pathway genes in cALL

To gain a better understanding of the underlying molecular mechanisms of how MondoA confers aggressiveness in cALL, we performed microarray analyses of various cALL cell lines (Affymetrix Human Gene 1.0 ST). For these experiments, we first compared Nalm6, 697 and REH harboring shRNA mediated MondoA knockdown with control cells expressing a scrambled shRNA. Second, we analyzed Nalm6 cells with MKO, using Cas9 Nalm6 cells containing lentiviral Cas9 only as control. Analysis indicated 151 and 255 significantly up- and down-regulated genes, respectively, upon knockdown of MondoA. MKO cells differentially expressed 1657 genes, with 1087 being upregulated (FC≧1.4) and 570 down-regulated (FC≦0.7). Notably, 321 (11.3%) gene alterations overlapped in both MKD and MKO cells, and show similar patterns in network analyses (Figure S2C). Gene Set enrichment analyses (GSEA) showed a significant downregulation of Myc target genes (HALLMARKS_MYC_TARGETS_V1 & V2), as well as oxidative phosphorylation (HALLMARK_OXIDATIVE_PHOSPHORYLATION), ROS pathways (REACTIVE_OXYGEN_SPECIES_PATHWAY), DNA repair (HALLMARK_DNA_REPAIR) and fatty acid metabolism (HALLMARK_FATTY_ACID_METABOLISM) genes in MondoA competent cells. This suggests an essential role of MondoA in regulating metabolic pathways and downregulating energy producing cascades as well as ROS-generating processes (Figure 4A, B, C). The validation of microarray data by qRT-PCR is shown in Supplemental Figure (S2D, E).

**Figure 4.**
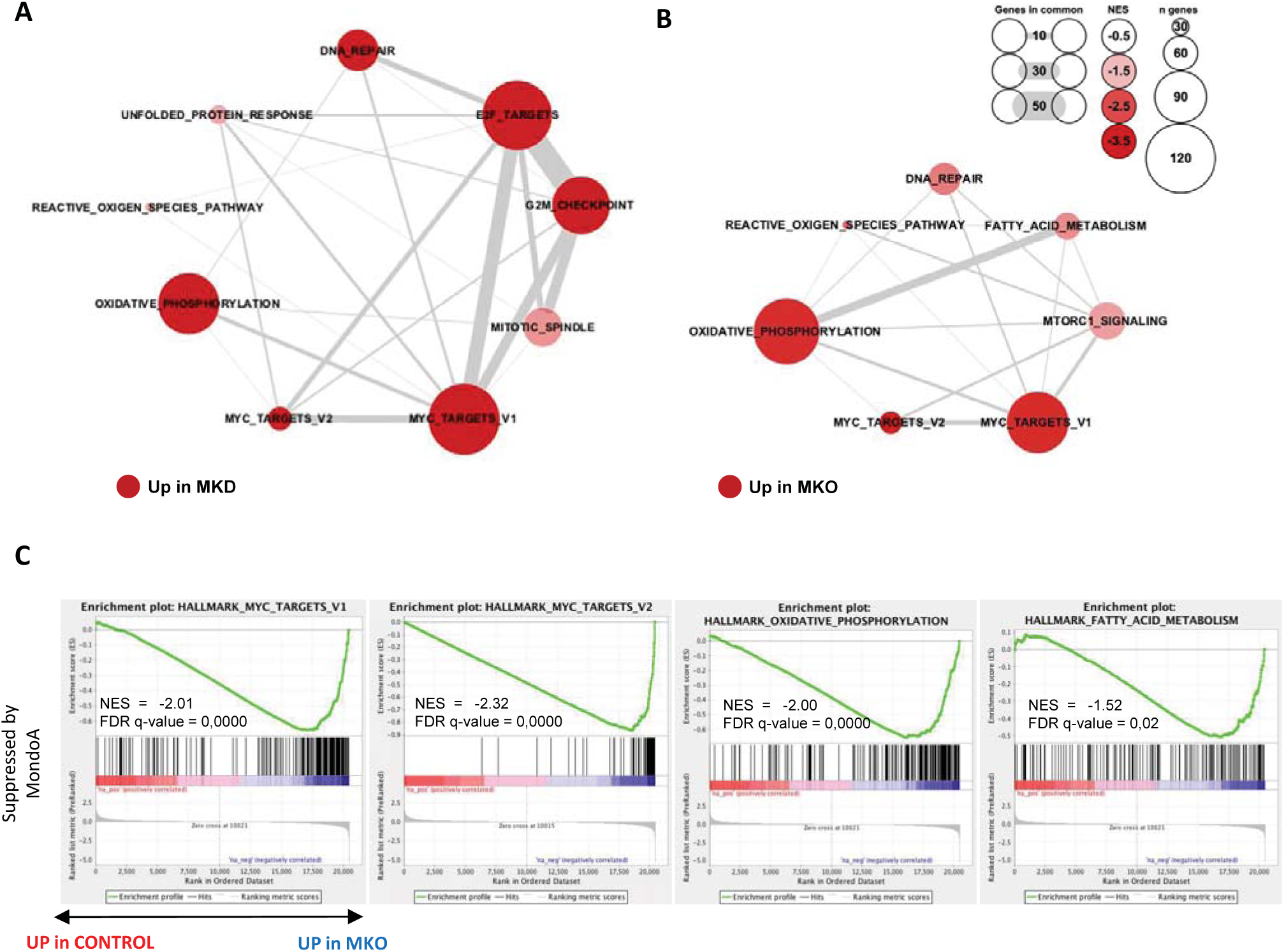
MondoA suppresses MYC-targets including fatty acid metabolism, oxidative phosphorylation and ROS pathway genes in cALL. **(A)** GSEA data summary represented as network generated using Cytoscape (version 3.7.1). Nodes depicted in the networks represent Hallmarks from GSEA with an FDR q-val < 0.05. Red color indicates negative NES (overexpressed in MKD (REH, Nalm6, 697) compared to control cell lines (REH, Nalm6, 697). Size of nodes corresponds to the number of genes in the pathway that belong to the core enrichment in gene set. Saturation intensity represents absolute NES. Edges connecting nodes represent number of genes that are shared between the two gene sets. **(B)** GSEA summary represented as network of Nalm6 cell lines either knockout for MondoA or control Nalm6. Data representation as described in (A). **(C)** Enrichment plots of significantly altered gene sets as yielded by the GSEA of microarray data of Nalm6 cell line with and without MKO. GSEA summary for MondoA knockout as well as GSEA summary for three cell lines (REH, Nalm6, 697) with knockdown in Supplementary (S3C, D). Gene sets stimulated by MondoA according to MKD and MKO experiments are shown in S3A, B. Heatmaps showing representative genes comprising MYC-target, OXPHOS, Fatty acid metabolism and Glycolysis gene sets are shown in S3E.

### MondoA provides leukemia stress-resistance by limiting oxidative phosphorylation via increased pyruvate dehydrogenase kinase (PDK) activity

To further investigate the metabolic restriction mediated by MondoA expression, we initially performed a proliferation assay comparing two MondoA deficient cell line clones with a clone overexpressing MondoA and control cells. MKO clones proliferated at significantly lower rates than both control and MondoA overexpressing cells (Figure 5A). Since our GSEA suggested that oxidative phosphorylation is reduced by MondoA, we measured mitochondrial respiration by determining cellular oxygen consumption rate (OCR) using an extracellular flux analyzer. MondoA deficient cells displayed a significantly higher rate of basal mitochondrial respiration compared to controls as well as cells overexpressing MondoA (Figure 5B, C). Additionally, ATP production (*i*.*e*. the difference between basal respiration and respiration following oligomycin treatment (Figure 5B, D)) and the maximum respiratory capacity (after addition of FCCP (Figure 5B, E)) were increased in MondoA deficient cells compared with controls. In line with this, overexpression of MondoA reduced oxidative phosphorylation back to control cell levels (Fig. 5B, C, D, E). Together, these findings indicate that MondoA limits oxidative phosphorylation (Oxphos) in cALL cells. It is known that pyruvate dehydrogenase (PDH) is an essential enzyme complex in the mitochondria that catalyzes the reaction from pyruvate to acetylCoA, which is subsequently utilized for the TCA cycle and Oxphos^26^. In view of the increased oxidative capacity of MondoA deficient cells, we assessed PDH activity in those cells, and found it was significantly increased compared to cells with MondoA overexpression or in vector alone control cells (Figure 5F). Control cells and cells with MondoA overexpression did not demonstrate significant difference in PDH activity. PDH is regulated through pyruvate dehydrogenase kinase (PDK), in which phosphorylation of PDH by PDK results in PDH inactivation^27^. Indeed, PDH showed reduced phosphorylation in MondoA deficient cells and conversely more phosphorylation in cells overexpressing MondoA (Figure 5G). Correspondingly, reduced PDK levels were observed at the protein and mRNA levels in the absence of MondoA (Figure 5G, H). Consistent with this, we observed a similar correlation in patient RNA-seq data sets (UCSC Xena, TARGET), whereby MondoA overexpression was highly correlated with increased expression of PDK1 transcripts (Figure 5I). This suggests that MondoA transcriptionally regulates *PDK* mRNA expression to reduce oxidative phosphorylation in cALL cells.

**Figure 5.**
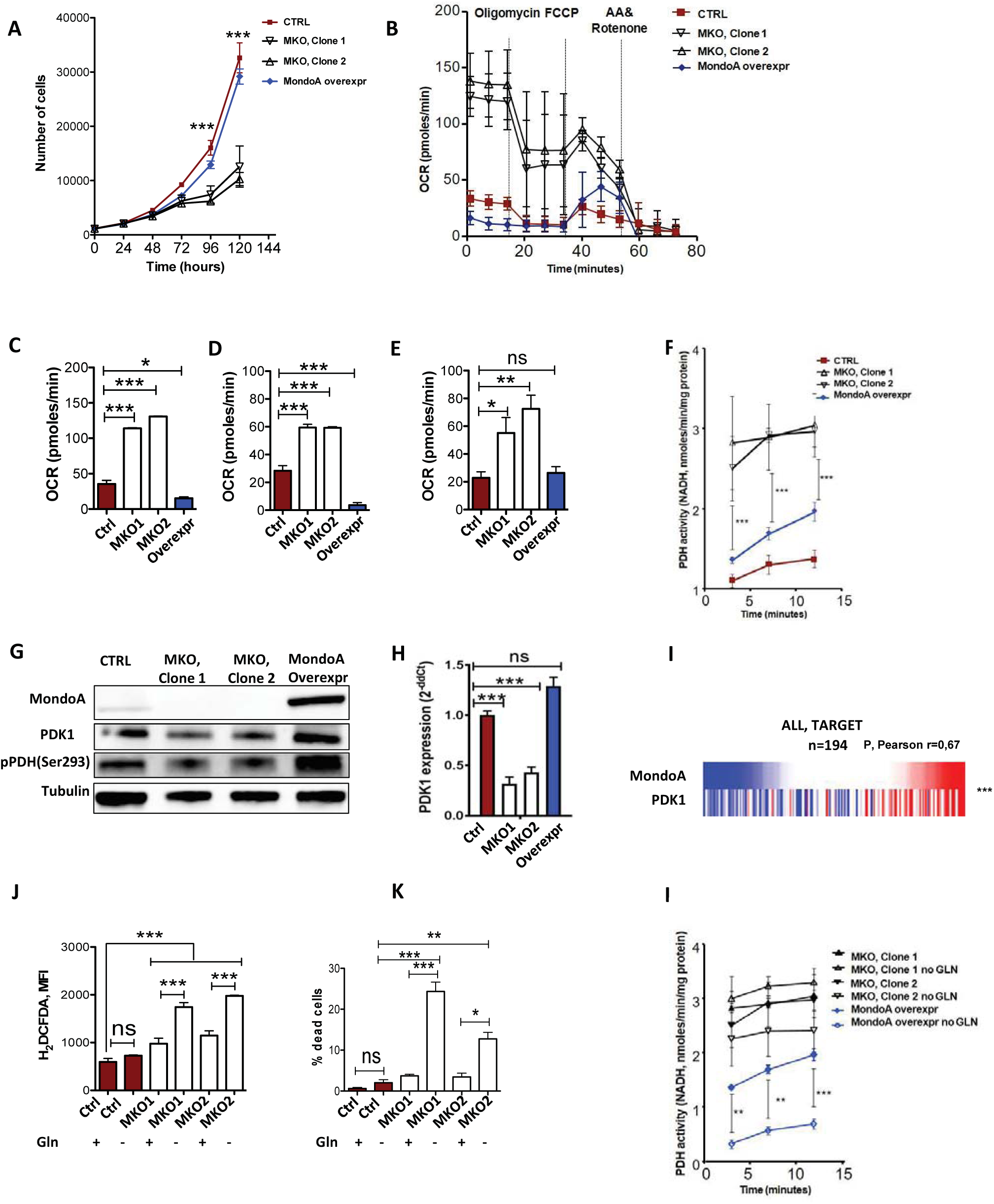
MondoA provides leukemia stress resistance by limiting oxidative phosphorylation and fatty acid synthesis via decreased PDH activity. **(A)** Long-term proliferation measured by direct cell counting. Control Cas9-only Nalm6 cALL cell line, MondoA overexpression and Nalm6 with MondoA knockout, grown under normoxia (20% O_2_) (n=6 for each time point, p<0.001 at 96h and 120h). **(B)** Mitochondrial respiration determined by cellular oxygen consumption rate (OCR) in control, MKO Nalm6 clones and Nalm6 cells with MondoA overexpression using a Seahorse extracellular flux analyzer (Agilent). **(C)** Basal respiration as measured OCR before Oligomycin injection (p<0.05 and p<0.001, 2way ANOVA with Bonferroni posttest). **(D)** ATP production as measured as the difference between basal respiration and OCR levels after Oligomycin injection. **(E)** Maximal respiration as measured by OCR levels after FCCP injection (p<0.05 and p<0.01, 2way ANOVA with Bonferroni posttest). **(F)** PDH activity assessed by NADH production. PDH activity in control, MKO and MondoA overexpressing Nalm6 cells assessed by colorimetric PDH activity assay kit (n=3 for each time point, p<0.001, 2way ANOVA with Bonferroni posttest). **(G)** Western blot showing MondoA, total PDK1 (Pyruvate Dehydrogenase Kinase 1), pPDH (Ser293) and alpha-Tubulin as loading control. MondoA overexpressing clone in comparison to MKO Clone 1 and 2 and Cas9-only control Nalm6 cells. **(H)** PDK1 mRNA in MKO clones compared to CTRL and MondoA overexpressing clones measured by qRT-PCR (p<0.001, 2way ANOVA with Bonferroni posttest). **(I)** Heat map showing PDK1 gene co-expression with MondoA in cALL. RNAseq data of 194 primary ALL patient’s samples (TARGET) from Xena browser. **(J)** Cellular ROS measurement of Nalm6 MKO and control cells under either GLN withdrawal or full media. H_2_DCFDA fluorescence measured by flow cytometry (n=3 for each experimental condition, p<0.001, one-way ANOVA with Bonferroni’s posttest) **(K)** Cell death rate determined by AnnexinV and 7AAD staining in control and MKO cells in either full medium or medium lacking glutamine (GLN) (n=3 for each experimental condition, p<0.05, p<0.01 or p<0.001, one-way ANOVA with Bonferroni’s posttest). **(L)** PDH activity assessed by NADH production. PDH activity in control, MKO and MondoA overexpressing Nalm6 cells with or without GLN starvation assessed by colorimetric PDH activity assay kit (n=3 for each time point, p<0.001, 2way ANOVA with Bonferroni’s posttest).

### MondoA confers adaptation to metabolic stress and limits ROS-production in cALL cells

Although the process of oxidative phosphorylation is highly efficient for generating ATP in cells, this process also poses high levels of endogenous ROS through the electron transport chain, which in turn can result in damage to DNA and proteins^28^. To investigate whether MondoA reduces oxidative phosphorylation in cALL cells to limit the production of ROS we measured ROS levels in the presence and absence of MondoA. Indeed, under normal media conditions control cells expressing MondoA generated significantly less ROS than MondoA deficient cells (Figure 5J). It is known that cells downregulate their TCA cycle activity under glutamine starvation, possibly to limit ROS accumulation^29-32^. To investigate whether MondoA similarly facilitates adaptation to metabolic stress, we conditioned cALL cells in glutamine-depleted culture medium for 24h. Notably, MKO cells showed higher ROS levels under glutamine starvation (Figure 5J), while no differences in ROS levels were observed in control cells under the same conditions. Moreover, while there was only a small increase in cell death in MKO compared to control cells under glutamine replete conditions, MKO cells showed a marked decrease in viability when starved of glutamine (Figure 5K). Notably, MondoA depletion resulted in an inability to reduce PDH activity under glutamine starvation; in contrast, cells overexpressing MondoA significantly reduced PDH activity under the same conditions (Figure 5L). Although further studies are necessary to fully uncover the mechanism, these data suggest that loss of MondoA renders cALL cells unable to reduce PDH activity and therefore the TCA cycle in response to starvation. We postulate that in this way, MondoA reduces ROS generation under nutrient stress conditions to protect proliferating cells from potentially lethal ROS accumulation.

## Discussion

Our findings establish MondoA as a factor that enables malignancy in common B-precursor ALL by providing adaptation to metabolic stress by dialing down Krebs cycle activity and the subsequent generation of reactive oxygen species. This is in line with our previous findings that MondoA promotes stemness, proliferation and B cell receptor signaling pathways^19^. Our findings support the notion that in glutamine dependent tumor entities, such as neuroblastoma or cALL, MondoA propagates malignancy^16^. However, in neuroblastoma the loss of MondoA results in down-regulation of oxidative phosphorylation and metabolic activity^15^. In cALL we see a different mechanism, whereby MondoA senses metabolic stress and restricts oxidative phosphorylation to a cell sustainable level by reducing PDH activity in glutamine scarce environments. Vice versa, MondoA loss enforces oxidative phosphorylation and PDH activity, which in turn cannot be modulated upon glutamine starvation. The process of oxidative phosphorylation is a major endogenous source of ROS and can result in cell lethality through oxidation of many essential components in a proliferating cell^33^. A carefully regulated ROS generation as well as scavenge mediated through MondoA is hence essential for sustaining malignant properties in cALL.

Additionally, we show the clinical relevance of high MondoA expression in cALL patient data sets. These data sets allowed us to explore the role of MondoA in cALL by assessing its high expression in cALL compared to solid tumors and other hematopoietic malignancies, and documenting its role for the aggressiveness of leukemia *in vitro* and *in vivo* as well as its consequences for gene expression and metabolic function. However, this remains a surprising finding since it has been described for glycolytic tumors, triple negative breast cancer or malignant melanoma, that MondoA downregulation is required for proliferation^18^. As mentioned earlier, in neuroblastoma an opposing role of MondoA has been described^16^.

In conclusion, our findings indicate that the role of MondoA in regulating metabolic processes is highly context specific and heavily depends on the metabolic profile of the specific tumor type. In pediatric cALL MondoA overexpression correlates with a worse clinical outcome due to the metabolic adaptive capabilities of MondoA. Interference with MondoA or its downstream targets, such as TXNIP, by candidate inhibitors^34^ could render those cells inept to adaption and hence be a novel therapeutic target for cALL.

## Acknowledgments

A.S. received grants from CURA PLACIDA, Children’s Cancer Research Foundation (CP102/120815) and TRANSAID Stiftung für Krebskranke Kinder (8810001358 GCP). Experimental support by Oxana Schmidt, Lynette Henkel is gratefully acknowledged. E.H. received a grant from the Else-Kröner-Stiftung. B.X. was supported by Chinese Scholarship Council (CSC: No. 201908210290). Laboratory of T.G.P.G. is supported by grants from the German Cancer Aid (DKH-70112257), the Gert and Susanna Mayer Foundation and the Barbara and Wilfried Mohr Foundation. M.B. was supported by Doris Stiftung. We also thank the following clinicians for providing samples used in this work: Irene Teichert - von Lüttichau, Angela Wawer, Katja Gall.

## Authorship Contributions

A.S. and E.H. coordinated and designed the study, performed all functional experiments, performed data analyses, wrote the paper, designed the figures and helped with grant applications. B.X. designed and performed most experiments required for resubmission. C.P. and M.B. did bioinformatic analysis and representation of microarray data. A.P. and A.G. provided guidance with ROS and Hypoxia experiments and data analysis. D.G.F. and D.B. helped to investigate metabolism. T.G.P.G. participated in the study design, provided compilation of independent published microarray studies, helped writing the paper and designing the figures, provided statistical guidance. J.S. analyzed the expression data of the BFM cohort patient’s samples. R.A.R. carried histologic analyses for in vivo experiment. M.C.B. carried out analyses of independent published microarray studies. G.C. provided expression data of the patients comprised the BFM cohort. G.R. provided biological and genetic guidance and helped with grant applications. J.R. supervised the study and provided laboratory infrastructure. P.S. supervised the study and helped writing the paper. S.B. initiated, designed and supervised the study; analyzed the data; wrote the paper together with A.S. and E.H. and provided laboratory infrastructure and financial support. All authors read and approved the final manuscript.

## Conflict of Interest Disclosures

SB has an ownership interest in PDL BioPharma, and served as consultant to EOS Biotechnology Inc., Bayer AG and Swedish Orphan Biovitrum AB. SB and GHR hold US and EU intellectual properties in gene expression analysis. Other authors have no conflict of interest to declare.

## Notes

### Competing Interest Statement

The authors have declared no competing interest.

